# Dynamics of bleomycin-induced lung injury at single-nucleus resolution

**DOI:** 10.64898/2026.09.08.750170

**Authors:** Cecilia Lopez-Martinez, Yu-Hua Chow, William A. Altemeier, Sina A. Gharib, Chi F. Hung

## Abstract

Single-cell RNA sequencing (scRNA-seq) has emerged as a powerful tool for studying individual cell populations in complex tissues during injury and repair. However, it involves extensive sample manipulation, limiting its application in rare and difficult-to-dissociate cell populations. Single-nucleus RNA sequencing (snRNA-seq) bypasses several of these challenges, but its performance in lung disease models is not well characterized. In a longitudinal murine model of bleomycin-induced lung injury, we performed whole lung snRNA-seq to systematically map 46 cell populations and states with minimal tissue processing. Our approach captured the emergence of Krt8+/Cdkn1a+ transitional alveolar cells and an expansion of macrophage populations during injury. Additionally, we identified sizeable populations of rare cell types, such as pericytes, and cells sensitive to digestion protocols, including a population of activated fibroblasts. These results indicate that snRNA-seq recapitulates the dynamic changes associated with injury as previously reported using single-cell methods and outperforms these approaches in the representation of cell types that are sensitive to processing, thereby highlighting its utility for high resolution analysis of heterogenous tissues such as lung.

## INTRODUCTION

Single-cell RNA sequencing (scRNA-seq) techniques have transformed our understanding of the lung as an integrated organ, allowing the identification of novel cell types in the steady state (1, 2), or the definition of dynamic cell states during injury and repair (3, 4). However, several technical limitations restrict its applicability. One key challenge is the need of enzymatic digestion of fresh tissue samples to obtain a single-cell suspension, which limits the use of this approach on tissues that are easily disaggregated, introduces bias against fragile cell types, and perturbs transcriptional activity. Also, the requirement for fresh tissue complicates the use of clinical samples and excludes archived frozen material.

Single-nucleus RNA sequencing (snRNA-seq) has emerged as a promising alternative that overcomes some of these limitations, as it requires minimal tissue processing and no enzymatic digestion. Indeed, a previous report has shown that both techniques are similar during homeostatic conditions (5). However, the capabilities and limitations of snRNA-seq haven’t been explored in the context of lung injury, where cell states undergo dynamic transcriptional changes and the potential limitations of missing cytoplasmic RNA in this approach remains unknown.

Here, we systematically assessed the fidelity of snRNA-seq to temporally profile transcriptomic changes at the cellular level during bleomycin-induced lung injury. We show that a single-nucleus approach without enzymatic digestion of lung tissue captures a broad range of lung cells, including rare cell types and cells sensitive to enzymatic digestion, while robustly recapitulating the transcriptional dynamics previously reported in the literature.

## MATERIALS AND METHODS

### Mouse model

Animal protocol was approved by the Institutional Animal Care and Use Committee at the University of Washington. Transgenic mice with tamoxifen-induced tdTomato expression in platelet-derived growth factor receptor beta (PDGFRb) lineage cells *(Pdgfrb*-Cre^ERT2/+^;*Rosa26*-tdTomato^flox/flox^) were provided a tamoxifen diet between PN weeks 3-7. Following 2 weeks of tamoxifen washout with normal chow, mice underwent bleomycin-induced lung injury as previously described (6–9). Detailed methods are available in the online data supplement. Mice (n = 2 per time point) were challenged with intratracheal bleomycin (1.3 U/kg) and lungs were harvested at days 7, 14 and 21 after injury. Lungs harvested at day 0 served as uninjured controls.

### Single-nucleus RNA-seq data generation

Whole lungs were flash-frozen and transported on dry ice to the Brotman Baty Institute Advanced Technology Lab (BAT-Lab) at the University of Washington for isolation of nuclei and library preparation using the sci-RNA-seq3 protocol (10, 11). Detailed methods are provided in the online data supplement.

### Single-nucleus dataset analysis

Removal of background RNA reads was performed using SoupX v1.6.2 (12) with rho = 0.25. Nuclei with less than 100 or more than 2500 genes profiled were filtered out. Dimensionality reduction and clustering were performed using Monocle3 v1.3.7 (10, 13–15). Clusters with consistently high Scrublet (16) scores and co-expression of markers from different cell types were identified as doublets and removed from the dataset. Annotation was performed using both canonical marker genes and previously published single-nucleus and single-cell RNA sequencing datasets (3, 5, 17).

### Whole lung single-nucleus (SN) vs. single-cell (SC) comparison

An equivalent dataset that sequenced bleomycin-injured mouse lung using scRNA-seq was found in a public repository (GSE141259). Detailed methods can be found in the original publication (18) and the online data supplement.

For data integration, the SN dataset was downsampled to match the number of cells of the SC dataset by random sampling, only the same time points from the SC dataset were considered, and both datasets were merged using the reciprocal PCA (RPCA) method.

For expression profile comparison analysis, samples were summarized into *in silico* bulk samples. Differential expression analysis was performed between the day 7 and day 0 conditions for each technique individually using DESeq2 v1.46.0 and a Benjamini-Hochberg adjusted P-value < 0.05 for significance (19). Log_2_ Fold Changes for all genes and both techniques were visualized in a scatter plot, correlation between both was drawn and statistical significance calculated using the Pearson coefficient. Proximity between samples was visualized using a PCA. Arranged loadings from PC1 were used as a ranked list for a Gene Set Enrichment Analysis (GSEA) using Reactome gene sets from MSigDB v25.1.1 (20) and ClusterProfiler v4.14.6 (21). GSEA was also performed for day 7 day versus day 0 in the SN and the SC analyses individually, with an FDR < 0.05 used to designate significant enrichment.

### Data availability

Raw and processed data from the single nucleus dataset have been deposited in the NCBI Gene Expression Omnibus (GEO) under accession number GSE304152. All code used for the analysis is available in https://github.com/cecilomar6/SCvsSN.

## RESULTS

### snRNA-seq captures lineage-specific changes associated with the dynamics of lung injury

The single-nucleus (SN) dataset profiled 24,324 genes in 103,836 nuclei from eight lung samples across four timepoints of bleomycin-induced lung injury. A total of 30 cell types and in 46 distinct states were identified by manual annotation (Figures 1A-B, Supplemental Table 1). Cell populations were categorized into lineages to study broad changes in composition during injury and repair (Figure 1C-D). The main differences across conditions were an expansion of the myeloid compartment at day 7 and a modest increase in mesenchymal cells. At days 14 and 21 alveolar epithelial cells increased while myeloid cells returned to baseline levels.

**Figure 1:**
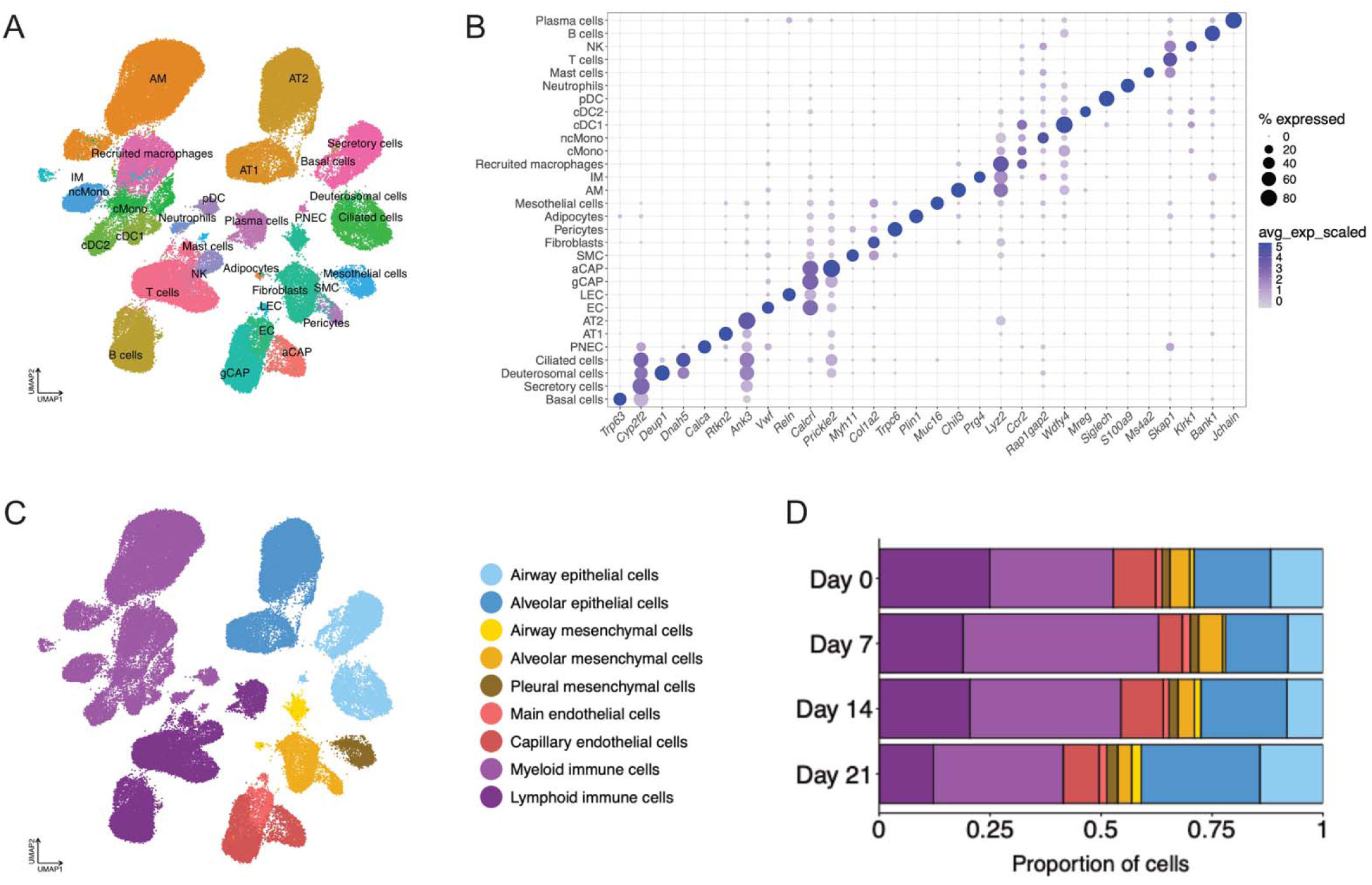
snRNA-seq captures cell population dynamics associated with lung injury. **A.** Uniform manifold approximation and projection (UMAP) plot showing 30 cell types identified in the SN dataset. AM = alveolar macrophages; AT1 = alveolar type 1 epithelial cells; AT2 = alveolar type 2 epithelial cells; EC = endothelial cells; IM = interstitial macrophages; LEC = lymphatic endothelial cells; NK = natural killer cells; PNEC = pulmonary neuroendocrine cells; SMC = smooth muscle cells; aCAP = alveolar capillary cells; gCAP = general capillary cells; cDC = conventional dendritic cells; pDC = plasmacytoid dendritic cells; cMono = classical monocytes; ncMono = non-classical monocytes. **B.** Dot plot showing the expression of canonical markers used for cell annotation for each cell type. **C**. UMAP showing the classification of each cell population into cell lineages. **D.** Bar plot showing the proportion of each cell lineage on each timepoint.

### snRNA-seq captures cell states associated with bleomycin injury in the epithelial, endothelial and immune compartments in whole lungs

We next performed a detailed characterization of each cell lineage.

In the epithelial compartment we identified 11 cell types and states, including some difficult to profile populations such as pulmonary neuroendocrine cells (PNEC) and basal cells (Figure 2A, Supplemental table 2). We also identified a damage associated transient progenitors-like cells, characterized by the expression of senescence marker *Cdkn1a*. This population was not present at baseline, appeared on day 7 and persisted through day 21 (Figure 2B).

**Figure 2:**
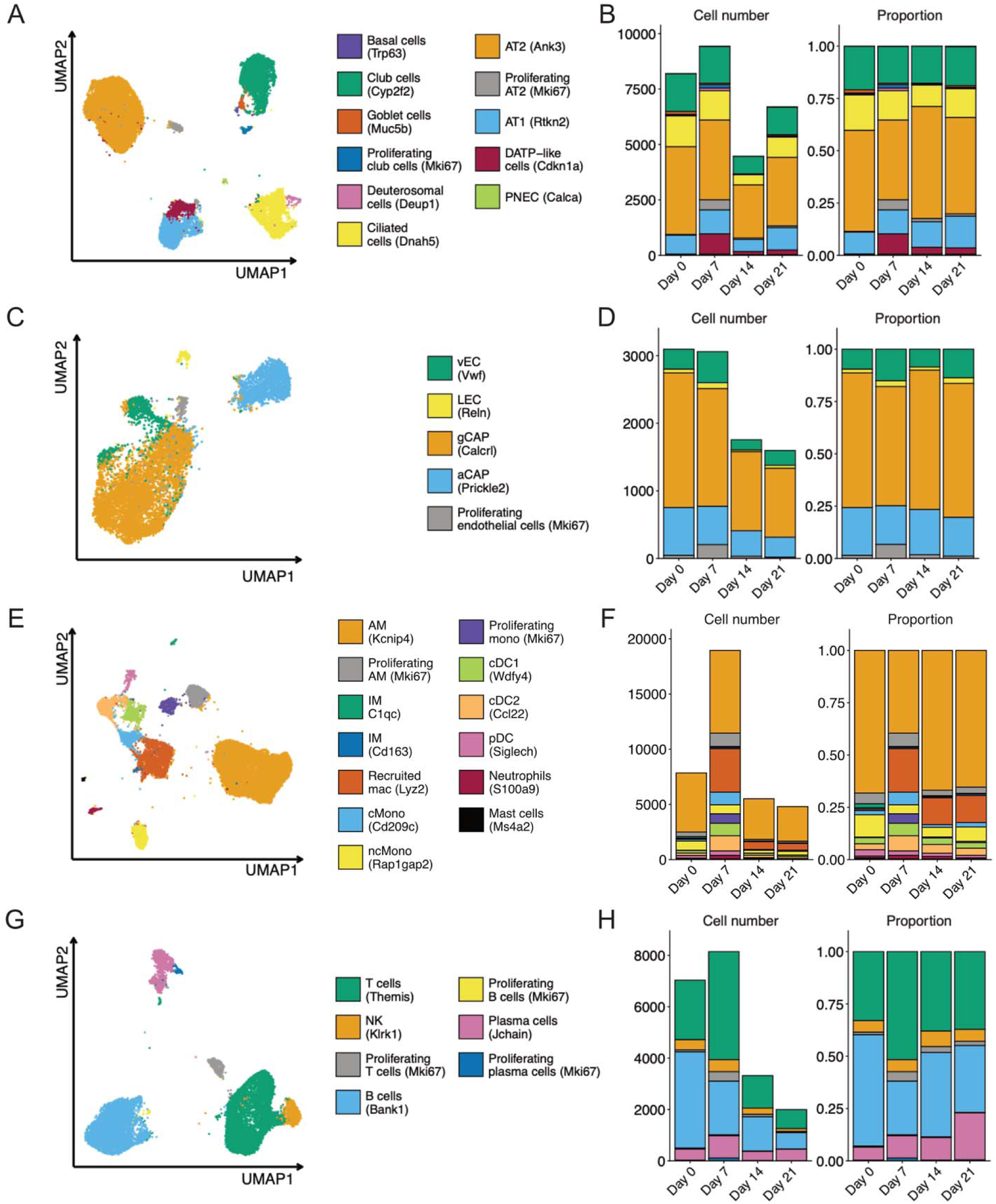
cell states identified and time dynamics for each lineage. **A, C, E and G.** Uniform manifold approximation and projection (UMAP) plot showing the cell states identified for the epithelial (A), endothelial (C), myeloid (E) and lymphoid (G) lineages. **B, D, F and H.** Bar plots showing the absolute cell numbers and relative proportion of each cell state for each timepoint, for the epithelial (B), endothelial (D), myeloid (F) and lymphoid (H) cell lineages.

We identified 5 cell populations in endothelial compartment, including both capillary, venous and lymphatic endothelial cells (Figure 2C, Supplemental table 3). We also identified a proliferative cell state that peaked at day 7 post-injury (Figure 2D).

Due to the complexity and dynamism of the immune lineage, we split this compartment into myeloid and lymphoid lineages. The myeloid compartment showed the biggest expansion after bleomycin injury. We identified 13 cell types and states within this compartment (Figure 2E, Supplemental table 4). The increase in cell number seemed to come from the appearance of multiple monocyte and monocyte-derived populations, including classical monocytes (cMono) and recruited macrophages (Figure 2F). We also observed an increase in proliferative populations, from both resident and recruited macrophages. In terms of the resolution of the technique, we were able to identify the presence of neutrophils and mast cells, which are usually difficult to profile due to high enzyme content in their cytoplasms.

We identified 7 cell types and states in the lymphoid lineage, including T cells, NK cells, B cells and plasma cells (Figure 2G, Supplemental table 5). We also identified proliferating populations for each of these cell types (Figure 2H).

### snRNA-seq identifies adventitial fibroblasts as participants in the injury response within the mesenchymal compartment

Single nucleus RNA-seq showed a great capacity to capture mesenchymal cells, allowing identification of rare mesenchymal cell populations such as pericytes and mesothelial cells (Figure 3A). The fibroblast cluster showed a marked expansion after injury (Figure 3B), prompting further exploration. Within this cluster, we identified five distinct cell subpobulations: alveolar and adventitial fibroblasts, two injury-associated populations and a proliferating population (Figure 3C, Supplemental table 6). Both the injury-associated populations and the proliferating population expanded after injury (Figure 3D).

**Figure 3:**
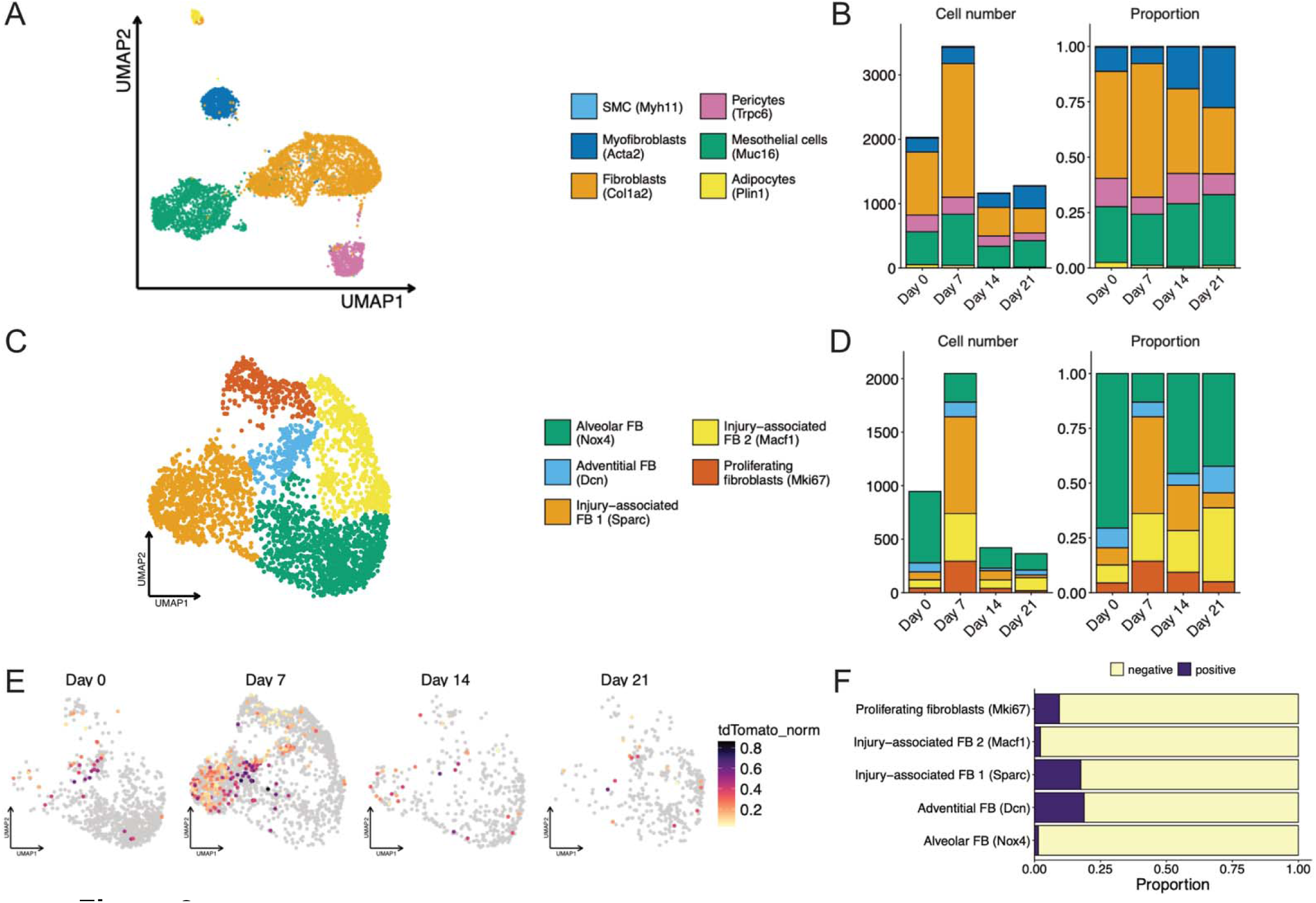
cell states and time dynamics of the mesenchymal compartment. **A.** Uniform manifold approximation and projection (UMAP) plot showing the cell types identified for the mesenchymal lineage. **B.** Bar plots showing the absolute cell numbers and relative proportion of each cell type for each timepoint. **C.** UMAP plot showing the distinct cell populations and states identified for the fibroblast cell population. **D.** Bar plots showing the absolute cell numbers and relative proportion of each fibroblast cell population for each timepoint. **E.** UMAP plots showing the tdTomato normalized expression in the fibroblast population, split by timepoint. **F.** Bar plot showing the percentage of tdTomato-positive cells for each fibroblast population.

In our transgenic lineage-tracing mice *Pdgfrb-Cre^ERT2^;Rosa26-tdTomato*, where tdTomato labeling occurred at baseline prior to bleomycin injury, tdTomato expression was restricted to Pdgfrb+ populations at baseline and their daughter cells. At baseline, tdTomato signal within the fibroblast cluster was restricted to the adventitial fibroblast population. By day 7, however, the predominant tdTomato+ population shifted to an injury-associated fibroblast state (Figures 3E and F). Nearly 25% of both adventitial and injury-associated 1 fibroblasts expressed tdTomato, suggesting that adventitial fibroblasts contribute to the fibrotic response elicited by bleomycin.

### scRNA-seq and snRNA-seq identify similar changes in cellular profiles during lung injury despite differences in gene expressions

To assess the capability of snRNA-seq to capture transcriptomic changes associated with lung injury and fibrosis, we compared it to a whole-lung single-cell (SC) dataset using an equivalent model. The SC dataset comprised 21,083 cells from 20 samples, profiling the expression of 23,400 genes. On average, SC cells contained 799 UMIs and 450 detected genes, while SN nuclei had 524 UMIs and 374 genes. A total of 38 cell populations had been previously annotated in the SC dataset.

After integration, cells from both techniques overlapped by lineage, indicating similar gene expression profiles (Figure 4A and B). As expected, immune cell populations expanded following lung injury in the SC samples. Similarly, SN samples showed an expansion of immune populations following injury, demonstrating both methods concordantly captured recruitment of immune populations in the lung during injury (Figure 4C)

**Figure 4.**
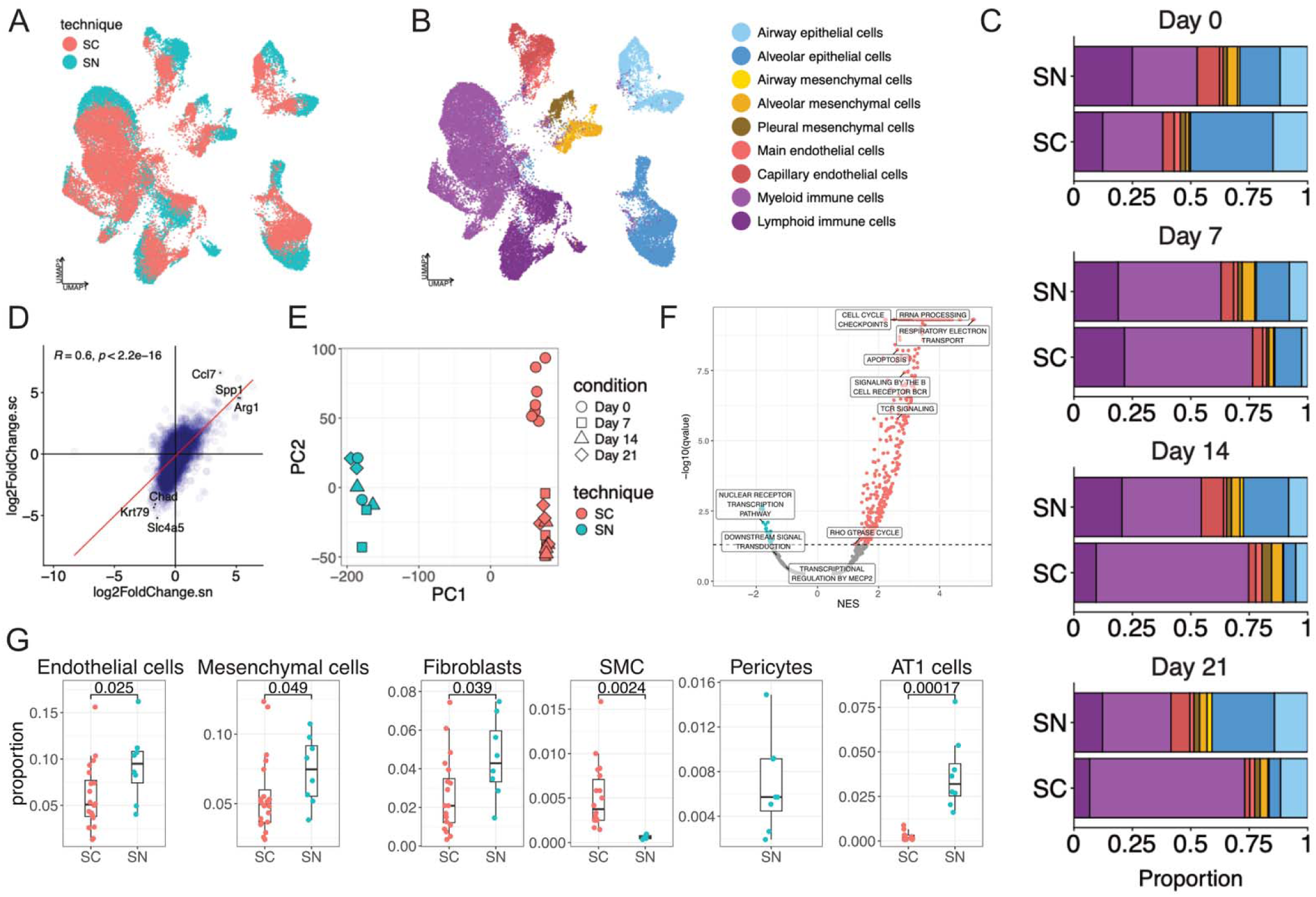
Differences and similarities between single-cell and single-nucleus profiling of whole lung. **A, B.** Uniform manifold approximation and projection (UMAP) plot showing the overlap between both datasets after alignment, colored by technique (A) and cell lineage (B). **C.** Bar plot showing the proportion of each cell lineage for each technique and timepoint. **D.** Scatter plot illustrating the correlation of the log_2_ Fold Change values between day 7 and day 0 for the SC and SN datasets. The red line represents the regression line. Statistical significance was assessed using Pearson correlation. Representative genes were labeled in the plot. **E.** Plot showing the 2 first principal components for the *in silico* bulk SC and SN samples. We can observe that PC1 separates the samples by technique. **F**. Gene weights from PC1 were used as a ranked list to perform a gene set enrichment analysis (GSEA). Volcano plot visualizes all gene sets tested, highlighting those significantly enriched in the SC (positive NES score), and SN (negative NES score) samples. Pathways considered relevant were labeled in the plot. **G.** Box plots depicting per-sample proportions of relevant cell populations, comparing SC and SN datasets. Statistical significance was assessed using a Wilcoxon test.

To study the similarity of the overall transcriptional response, we performed differential gene expression analysis between day 7 vs. day 0 for each technique using *in silico* bulk samples (Supplemental Tables 7 and 8). The resulting transcriptional profiles showed a positive correlation (R = 0.55, p < 2.2 x 10^−16^, Figure 4D), indicating similar global expression changes.

We then performed gene set enrichment analysis (GSEA) to compare the ability of SN and SC to detect deregulated pathways during bleomycin lung injury. Both techniques identified an upregulation of pathways related to antigen presentation and myeloid mediated immunity, and a downregulation in cilium motility. SC uniquely identified an upregulation in pathways related to B cell activity, and SN an upregulation in processes related to cell division. There were also some discordant results between the two techniques: while SN detected an upregulation of cell differentiation and metabolic processes, SC identified the same pathways as downregulated (Supplemental Table 9). We also performed GSEA based on the weights of the principal component that separated the samples by technique. We found a higher expression of genes related to ribosomal, mitochondrial and immune cell receptor activity in SC samples, while SN samples were enriched in genes related to chromatin remodeling and transcription factor activity (Figures 4E and F, Supplemental Table 10).

Cells with an elongated morphology or strong attachments to the extracellular matrix, such as endothelial and mesenchymal cells, or in the specific case of the lung, AT1 cells, are known to be challenging to profile using scRNA-seq. Our SN dataset showed consistently a significant higher proportion of endothelial cells, mesenchymal cells and AT1 cells (Figure 4G). Additionally, we found notable differences in the profiling of specific mesenchymal populations, with SN having significantly higher proportions of fibroblasts and uniquely identifying a population of pericytes, but underperforming in the identification of smooth muscle cells.

Overall, these results indicate that snRNA-seq captured similar changes in cellular populations and transcriptional activity associated with lung injury than scRNA-seq. Despite differences in tissue processing and analysis methodologies, the SN protocol was able to capture dynamic changes associated with lung injury that were previously characterized using more widely used SC methods, while also capturing a broader representation of many key lung cell populations.

## DISCUSSION

Profiling of tissues at the single-cell level is a rapidly evolving field, especially in the areas of injury and fibrosis, but the adoption of novel techniques requires evaluation for efficacy and applicability. In real world application of RNA sequencing methodologies, animal use and cost constraints often limit investigators to the use of one type of single-cell technology. With the single-nucleus approach, the question we ask is whether it captures dynamic changes in cell states and transcriptional responses similar to published experience using single cell methods. Here, we present the first temporal dataset of bleomycin-induced lung injury profiled using single-nucleus RNA sequencing, and compare it to equivalent, publicly available, single-cell RNA sequencing datasets.

Single-cell techniques preceded single-nucleus methods and have become standard for transcriptomic studies. snRNA-seq was initially developed to study brain tissue, which is challenging to dissociate due to the size, shape and intertwining of its cells. These studies demonstrated that RNA sequencing of isolated nuclei was technically feasible and provided adequate information to identify distinct cell types and states (10, 22–24). However, since it is not the standard approach, reference datasets for other organs and diseases remain limited.

In the lung, a previously published work compared single-cell and single-nucleus approaches in steady state, but similar comparisons during injury and repair have not been performed (5). Here, we show that nuclear RNA profiling can robustly capture these dynamic conditions, and for some vulnerable cell populations, improve upon cytoplasmic RNA sequencing. We found that both methods captured similar overall differential gene expression profiles in response to injury. The main differences in the transcriptional signal was an enrichment of genes related to cytoplasmic processes in the SC dataset whereas SN was enriched in genes involved in nuclear processes. Of note, the SC dataset also showed an enrichment of genes related to immune cells receptors, and SN had reduced expression of canonical immune markers. This inability to capture key well-known immune-related transcripts highlights the need for further characterization of nuclear markers in immune-focused studies, and represents a potential limitation for SN-based analyses of specific immune populations.

The SC dataset used for the comparison was originally reported in a publication that identified a *Krt8*+ alveolar differentiation intermediate during alveolar regeneration. (18) We were able to identify a similar population in our SN dataset (DATP-like cells (*Cdkn1a*)), which confirms that SN profiling captures comparable repair cell states in the epithelial compartment as previously documented using SC approach and validates the utility of SN techniques to interrogate transcriptomic profiles during injury.

Cell types integral to lung biology such as AT1 cells and pericytes are strikingly underrepresented in single-cell datasets. One potential explanation for this observation is that these cell types are highly sensitive to enzymatic and mechanical disaggregation required by single-cell techniques. Particularly, pericytes are an understudied mesenchymal population with important regulatory roles in lung injury and repair (7, 9), but the difficulty of isolating and sequencing them has limited their characterization to date. In our SN dataset, however, we identified a sizeable pericyte population, indicating that this approach is better suited for studying this elusive cell type.

Another cell type that can be sensitive to the harsh processing of tissue disaggregation is endothelial cells. Morphometric analysis of the human lung reveals that endothelial cells constitute approximately 30% of the cells in the alveolar region (25) and up to 43% of lung parenchymal cells in rats (26). However, whole-lung SC datasets consistently display far lower frequencies of these cells than suggested by morphometry. Our SN dataset detected double the number of endothelial cells compared to SC, offering a much more robust representation of this key cell population in the lung.

Mesenchymal cells are the central effectors of matrix deposition and remodeling. Because of their embedding within the extracellular matrix, they are also notoriously difficult to profile in whole-tissue single-cell samples. Strategies to address this challenge often involve sample enrichment by sorting, which may alter *de novo* transcriptional states. We were able to identify a sizeable mesenchymal population without any sorting-based enrichment. The cellular origin of profibrotic fibroblasts during lung fibrosis remains debated, with some studies identifying the adventitial fibroblasts while others pointing to alveolar fibroblasts (4, 27). Our *Pdgfrb-Cre Rosa26-tdTomato* lineage-trading model shows tdTomato signal restricted to adventitial fibroblasts and injury-associated fibroblasts, indicating that adventitial fibroblasts, even if not the sole contributors, participate in the fibrotic process.

Our work has some limitations. The use of a cell suspension instead of nuclei was not the only technical difference between the SC and the SN datasets, since different protocols for the experimental model, library generation and sequencing platforms were used. We observed large differences in the number of cells/nuclei obtained per sample: the whole-lung SC dataset included approximately 21,000 cells from 20 samples, whereas the SN dataset comprised around 100,000 nuclei from eight samples. This stark difference is likely a consequence of the library preparation protocol used, as combinatorial indexing methods are optimized for their scalability to larger numbers of cells/nuclei (10). The larger number of cells may also explain the improved detection of rare cell types in the SN dataset. Also, bleomycin doses used were similar but not identical, which could have influenced the response to injury observed in each dataset.

In conclusion, we show that profiling of whole lung samples using snRNA-seq successfully recapitulates dynamic changes associated with bleomycin-induced lung injury previously reported using scRNA-seq and outperforms single-cell techniques in the representation of cell types sensitive to digestion protocols such as mesenchymal and endothelial cells. Given that snRNA-seq can be performed with minimal tissue processing, provides a more accurate representation of the lung’s cellular composition, allows the use of frozen samples, and facilitates sequencing of archived samples in a single run to minimize batch effect, it represents an attractive alternative to single-cell based methods for investigation of lung injury and fibrosis.

## AKNOWLEDGEMENTS

We thank the Brotman Baty Advanced Technology Lab (BAT-Lab) at the University of Washington for the sequencing services and expertise provided at the platform.

## Author contributions

CFH and SAG designed the research study. CFH and YHC conducted the experiments. CLM, CFH and SAG analyzed the data. CLM wrote the manuscript and prepared the figures. CFH, WAA and SAG contributed to editing and revising the manuscript.

## Artificial Intelligence disclosure

AI was used for proofreading of manuscript drafts. The authors reviewed, edited, and approved all editing suggestions. No AI tools were used to generate original content including data, figures, and the manuscript text.

